# K_V_1.2 channels inactivate through a mechanism similar to C-type inactivation

**DOI:** 10.1101/784249

**Authors:** Esteban Suárez-Delgado, Teriws G. Rangel-Sandín, Itzel G. Ishida, Gisela E. Rangel-Yescas, Tamara Rosenbaum, León D. Islas

## Abstract

C-type inactivation has been described in multiple voltage-gated K^+^ channels and in great detail in the *Drosophila Shaker* channel. As channels have moved into the structural era, atomic details of this and other gating mechanisms have started to be better understood. To date, the only voltage-gated channels whose structure has been solved are KvAP (X-ray diffraction), the K_V_1.2- K_V_2.1 “paddle” chimera (X-ray diffraction), K_V_1.2 (Cryo-EM); and ether-á-go-go (Cryo-EM) (Wang and MacKinnon, 2017), however, the characteristics and mechanisms of slow inactivation in these channels are unknown or poorly characterized. Here we present a detailed study of slow inactivation in the rat K_V_1.2 and show that it has some properties consistent with the C-type inactivation described in *Shaker*. We also study the effects of some mutations that are known to modulate C-type inactivation in *Shaker* and show that qualitative and quantitative differences exist in their functional effects, possibly underscoring subtle but important structural differences between the C-inactivated states in *Shaker* and K_V_1.2.

## Introduction

Voltage-dependent potassium channels undergo several types of gating processes that limit the conductance on a time-dependent manner. Among the many gating transitions of this type is a slow inactivation process termed C-type inactivation. Although it was described 28 years ago (Hoshi et al., 1991), this slow process is still not fully understood (Hoshi and Armstrong, 2013).

Inactivation processes in potassium channels have different functional manifestations and molecular origins (Hoshi et al., 1991; Kurata and Fedida, 2006). While rapid, ball and chain type inactivation is mediated by the N-terminus or β-subunits in *Shaker*-type channels (Hoshi et al., 1990; Rettig et al., 1994; Vergara-Jaque et al., 2019), slow inactivation is a more subtle process which likely involves conformational changes in the pore domain and the selectivity filter (De Biasi et al., 1993). Slow inactivation has been divided into C-, P- and/or U-types, as these might be unique or coexist in the same channel. Their different corresponding molecular mechanisms, if they exist, are not clear (Loots and Isacoff, 1998; Cordero-Morales et al., 2007; Cheng et al., 2011; Bähring et al., 2012b). In this paper, we will use the term slow inactivation, which in this case might be identified with the C-type inactivation amply described in *Drosophila Shaker* channels with genetically-removed fast inactivation (Hoshi et al., 1990; Klemic et al., 2001). K_V_1.2 are mammalian *Shaker*-like potassium channels that show up to 82% homology with the *Drosophila Shaker* gene product and have become an indispensable model for biophysical studies of potassium channels, principally because the availability of structures determined by X-ray diffraction as well as cryo-EM (Long et al., 2005; Pau et al., 2017; Matthies et al., 2018). These structures serve as templates to interpret biophysical data arising from other voltage-gated potassium channels. As mentioned, another important model in channel biophysics is the *Shaker* channel, especially mutants in which fast N-type inactivation has been removed. However, no high-resolution structures of *Shaker* exist and all experimental observations in *Shaker* are interpreted using the K_V_1.2 structures or computational models based on K_V_1.2. Recently a structure determined by cryo-EM was obtained for the K_V_1.2 channel in nanodiscs (Matthies et al., 2018) and it was suggested that the structure might possibly correspond to a C-type slow-inactivated state. However, no comprehensive description of slow inactivation of K_V_1.2 is available, although previous work has indicated the presence of a slow inactivation process as well as feasible similarities between *Shaker* C-type inactivation and slow inactivation in K_V_1.2 (Cordero-Morales et al., 2011). Our goal in this work was to characterize the inactivation process in rat K_V_1.2 channels and to investigate the effect of some equivalent mutants that in *Shaker* channels affect C-type inactivation. We find that K_V_1.2 channels are slower to inactivate than *Shaker* and that, although this might be identified with C-type inactivation, the effects of mutations and ions are sufficiently distinctive to suggest that the detailed slow inactivated conformation of K_V_1.2 is different from that of *Shaker*.

## Materials and Methods

### Channels, mutagenesis and oocyte injection

The rat K_V_1.2 channel in the pMAX plasmid was a gift courtesy of B. Roux (University of Chicago). Mutagenesis was a carried out by a single PCR reaction using the KOD Hot Start DNA polymerase (Novagen) and appropriate mutagenic oligonucleotides (Sigma). Mutants were confirmed first by restriction essays and finally by sequencing at the Molecular Biology Facility at the Instituto de Fisiologia Celular-UNAM. WT K_V_1.2 and the generated mutants’ cDNA where linearized with PacI endonuclease. Messenger RNA (mRNA) was synthetized from linearized cDNA with the T7 polymerase mMesage mMachine kit (Ambion) and resuspended in pure H_2_O at a concentration of 0.5-1 μg/μl.

*Xenopus laevis* handling was done according to NIH standards (National Research, 2010). Stage VI oocytes were obtained by surgery after frogs were anesthetized with tricaine (2.2 mg/ml). Oocyte defoliculation was achieved first by mechanical separation with fine tweezers and then by enzymatic treatment with collagenase type IA (Sigma-Aldrich) at a concentration of 1.2 mg/ml in calcium-free OR2 medium for 30 min. After thorough washing in calcium-free ND96 solution for 30 min, oocytes were kept in regular ND96 solution. Healthy oocytes were microinjected with ~40 nl of K_V_1.2 WT or mutant mRNA one day after surgery and experiments carried out between 2 and 6 days after injection. For patch-clamp recording, the viteline membrane was mechanically removed with fine tweezers before the experiments.

### Electrophysiology

#### Two-electrode voltage-clamp

Ionic and gating current recordings were performed using a two-electrode voltage-clamp (TEVC, Warner Instruments, OC-725C). Glass microelectrodes were fabricated in P-97 micropipette puller (Sutter Instruments) from borosilicate glass capillaries (BF150-86-10, Sutter Instruments) and had a resistance of 1-2 MΩ when filled with 3 M KCl and dipped in ND96 solution. Current was filtered at 1 kHz with an analog filter (Frequency Devices) and sampled at 10 kHz. Ionic currents where recoded in response to voltage-clamp pulses and using a –p/4 subtraction protocol to remove linear components of the current. Currents obtained from long pulses were not subtracted.

Gating currents were recorded in TEVC with ND96 as external solution. Linear current components were subtracted using a −p/4 subtraction protocol. Gating currents were sampled at 28 kHz. Ten sweeps were averaged at each voltage. These recordings were obtained from the double mutants: W366F/V381T, W366F/V381A, W366F/V381I. The mutants W366F/V381W, W366F/V381L did not show functional expression.

Voltage control and current recording were accomplished with the freely available program WinWCP running on a Windows XP PC (Dr. John Dempster, University of Strathclyde http://spider.science.strath.ac.uk/sipbs/page.php?page=software_ses) and a National Instruments (NI USB-6251) card. All recordings were carried out in ND96 as the bath solution unless otherwise indicated in the text.

#### Patch-clamp

Currents in the inside-out configuration of the patch-clamp were recorded with an Axopatch 200-B amplifier (Molecular Devices) and an Instrutech ITC-18 interface (HEKA Elektronik) controlled by PatchMaster software (HEKA Elektronik). Pipettes had resistance between 0.5-1 MΩ and were filled with a solution containing: 60 KCl, 60 NaCL, 3 HEPES, and 1.8 CaCl2, pH 7.4 (KOH). The bath solution contained: 130 KCl, 3 HEPES, and 1 EDTA, pH 7.4 (KOH). For non-stationary noise analysis, 50-300 current sweeps were recorded, filtering at 10 kHz and sampling at 50 kHz in response to a pulse from −90 mV to 100 mV with duration of 100 ms. No subtraction of linear current components was applied. The variance was calculated from the current sweeps using pairwise subtraction to reduce the influence of channel run-down or run-up (Heinemann and Conti, 1992). The variance vs. mean relationship was plotted and fit to the equation (Sigworth, 1980):

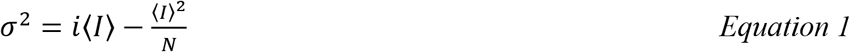

In this equation, *σ*^*2*^ is the variance of the current, *i* is the unitary channel current, *N* is the number of channels in the patch and ⟨*I*⟩ is the current average. The channel open probability, *P*_*o*_, was obtained from these parameters as: *P*_*o*_ = ⟨*I*⟩/*iN*.

#### Data analysis

Analysis was carried out with procedures written in-house in Igor Pro 6.0 (Wavemetrics Inc.). The conductance, *G(V)*, at each voltage *V* was measured from steady-state ionic currents, *I*, as: *G*= *I*/*(V-V*_*rev*_), were *V*_*rev*_ is the reversal potential. For inactivating currents, the peak current was measured to calculate the peak conductance. The normalized conductance was defined as: *G*/*G*_*max*_ were *G*_*max*_ is the conductance at 50 mV. *G*/*G*_*max*_ was fit to a Boltzmann-like function:

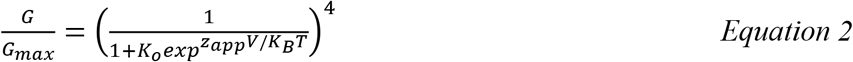

This equation assumes the sequential movement of four identical subunits with apparent charge movement per subunit given by z_app_. The parameter K_o_ describes the position of the curve along the V- axis. We find that this equation provides a better fit to the G-V curves than a simple Boltzmann function.

The slow ionic current decay elicited by long, 20 s pulses, was fit to the equation:

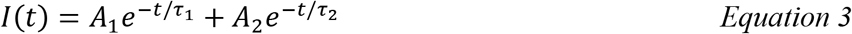

The A_i_ s are the amplitudes of the slow and fast components and the τ_i_ s are the respective time constants.

Steady-state inactivation was assessed with a conventional prepulse protocol. For WT K_V_1.2 the prepulse had a duration of 20 s and was changed from −80 mV to 70 mV in 10 mV increments. Then the voltage was brought to 50 mV during 300 ms. For the W366F mutant, the prepulse lasted 300 ms and was carried from −120 mV to 10 mV in 10 mV increments. After the prepulse the voltage was stepped to 50 mV for 100 ms.

The current at the 50 mV pulse was normalized and plotted as a function of prepulse voltage. These data were fit to the following equation.

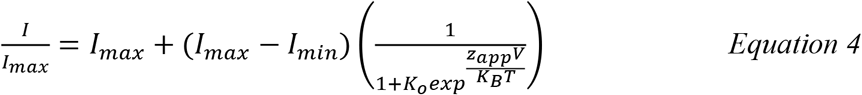

The ON-gating currents were integrated numerically to obtain the charge movement, *Q*_*on*_. *Q*_*on*_ was plotted as a function of voltage and fit to the equation:

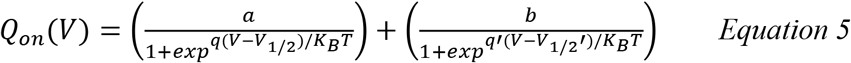

This equation assumes two main components of charge movement *q* and *q’*, with amplitudes *a* and *b* and mid-points of charge movement given by the values of V_1/2_ and V_1/2_’. The voltage dependence of any given time constant was fit to equation:

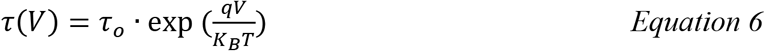

Here τ_o_ is the value of the time constant at 0 mV, *q* is the partial charge and *V*, *K*_*B*_ and *T* have the same meaning as in the previous equations.

#### Molecular modeling

Mutations were introduced in the K_V_1.2- K_V_2.1 paddle chimera channel (PDB: 2R9R, (Long et al., 2007)) using the mutagenesis module in PyMol. The rotamers with less molecular clashes were chosen to represent the final model. Structural figures were rendered in PyMol.

## Results

As previously shown (Roberds and Tamkun, 1991; Tao and MacKinnon, 2008; Ishida et al., 2015), K_V_1.2 produces steeply voltage-dependent, outward rectifying potassium currents (Fig. 1a and 1b). In the absence of co-expression with a β-subunit, these currents do not inactivate in a time scale of milliseconds, however, when long positive voltage pulses (20 s) are applied, the currents decay with a double-exponential time course with time constants of ~ 1 and 20 s at voltages more positive than 20 mV (Fig.1c and d). At 40 mV and after 20 s, 49.2 ± 0.21 % of the current remains (Fig. 1f). For comparison, the slow inactivation in *Shaker* channels with the N-terminal inactivation particle removed, is also bi-exponential with comparable time constants ~4 and 24 s, but the reduction of current at the end of the pulse is more complete (Olcese et al., 1997). At steady-state, the voltage dependence of inactivation, evaluated by a prepulse experiment, is very steep, with and apparent voltage dependence equivalent to ~5 e_o_ (Fig.1e and f). Inactivation is incomplete and shows a non-monotonic behavior reminiscent of U-type inactivation described in other potassium channels (Klemic et al., 1998).

**Figure 1.**
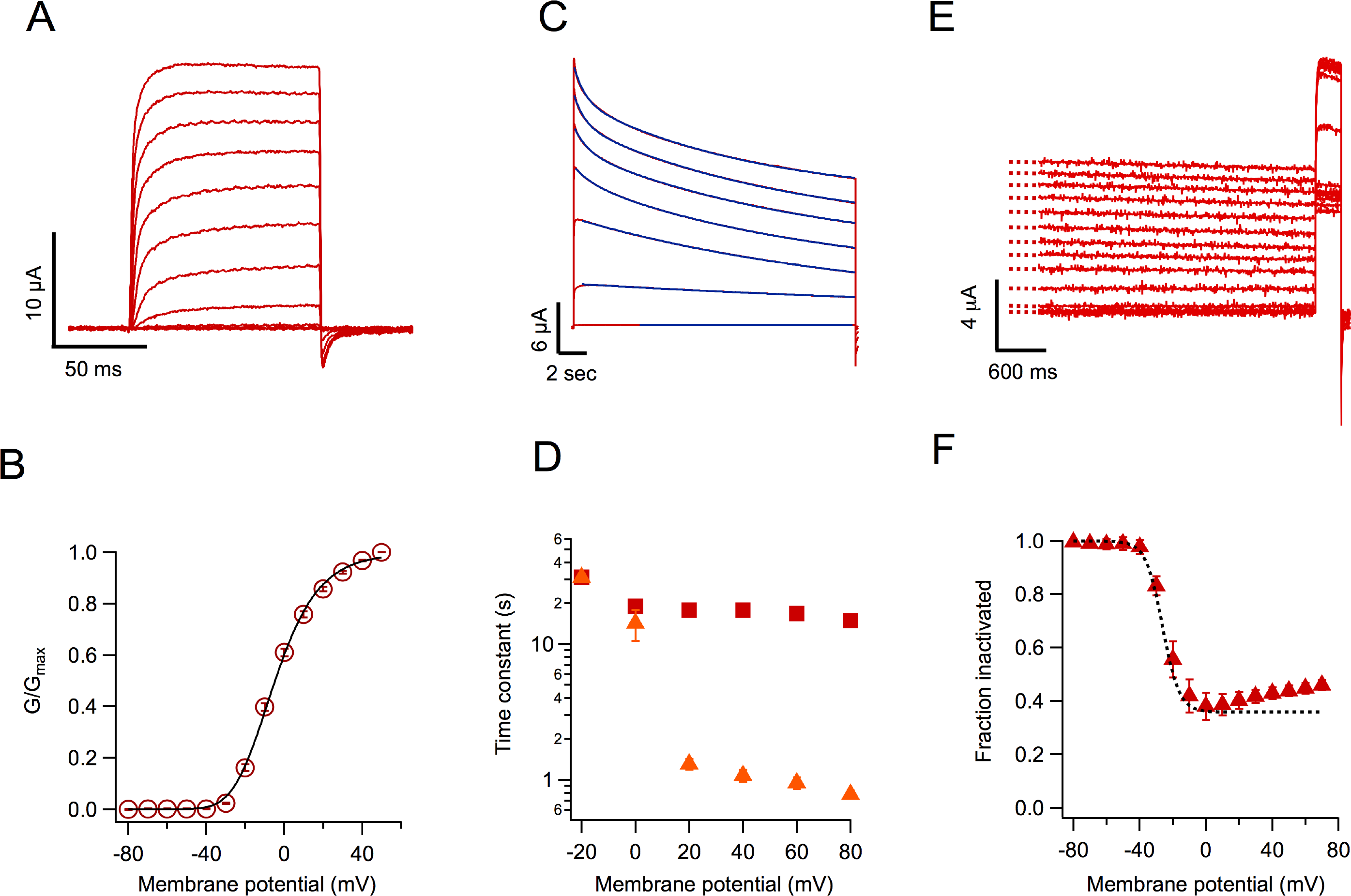
K_V_1.2 wild-type channels slow-inactivate incompletely. (A) Currents in response to short voltage-clamp pulses to 50 mV applied from a holding potential of −80 and lasting 100 ms. (B) Voltage–dependence of the conductance. The data are the mean and S.E.M. from n = 7 oocytes. The black curve is the fit of the data to Equation 2 with parameters: K_o_ = 0.13, z_app_ = 1.77 e_o_. (C) Currents in response to long (20 s) pulses from −40 mV to 80 mV in 20 mV steps. The current slowly decays in amplitude as a result of a slow inactivation process. The decay kinetics can be fit by a sum of two exponential functions. The fit to equation 3 is shown by the continuous blue lines. (D) Inactivation time constants are plotted as a function of voltage. The square symbols are the slow time constant and the triangles the fast time constant. (E) Protocol to determine the voltage dependence of steady state-inactivation at the end of a 20 s pulse. The current elicited by a 50 mV pulse and 300 ms duration after the long 20 s pulses is reduced as a consequence of accumulated inactivation. (F) Quantitation of the steady-state inactivation in an h∞-like curve. Notice that inactivation is relieved at voltages more positive than 20 mV. The dashed curve is the fit of a Boltzmann-like function, with an apparent valence of 5 e_o_. The difference at positive voltages between the fit and the data emphasizes the U-type inactivation character of the process.

An important hallmark of C-type inactivation in other channels is its modulation by extracellular cations, in particular potassium ions. Higher potassium extracellular concentrations ([K^+^]_o_) slowdown the inactivation rate (López-Barneo et al., 1993). We find that inactivation of K_V_1.2 channels is also slowed down by increased [K^+^]_o_ (Fig. 2a), although the effect is not very large. The rate of slow inactivation can also be modulated by extracellular tetra-ethyl-ammonium (TEA). Addition of 20 mM TEA to the bath decelerates inactivation by a similar extent to that of 100 mM external potassium (Fig. 2b). These behaviors with respect to external K^+^ and TEA, are hallmarks of C-type inactivation as seen in *Shaker* (Choi et al., 1991; López-Barneo et al., 1993; Andalib et al., 2004).

**Figure 2.**
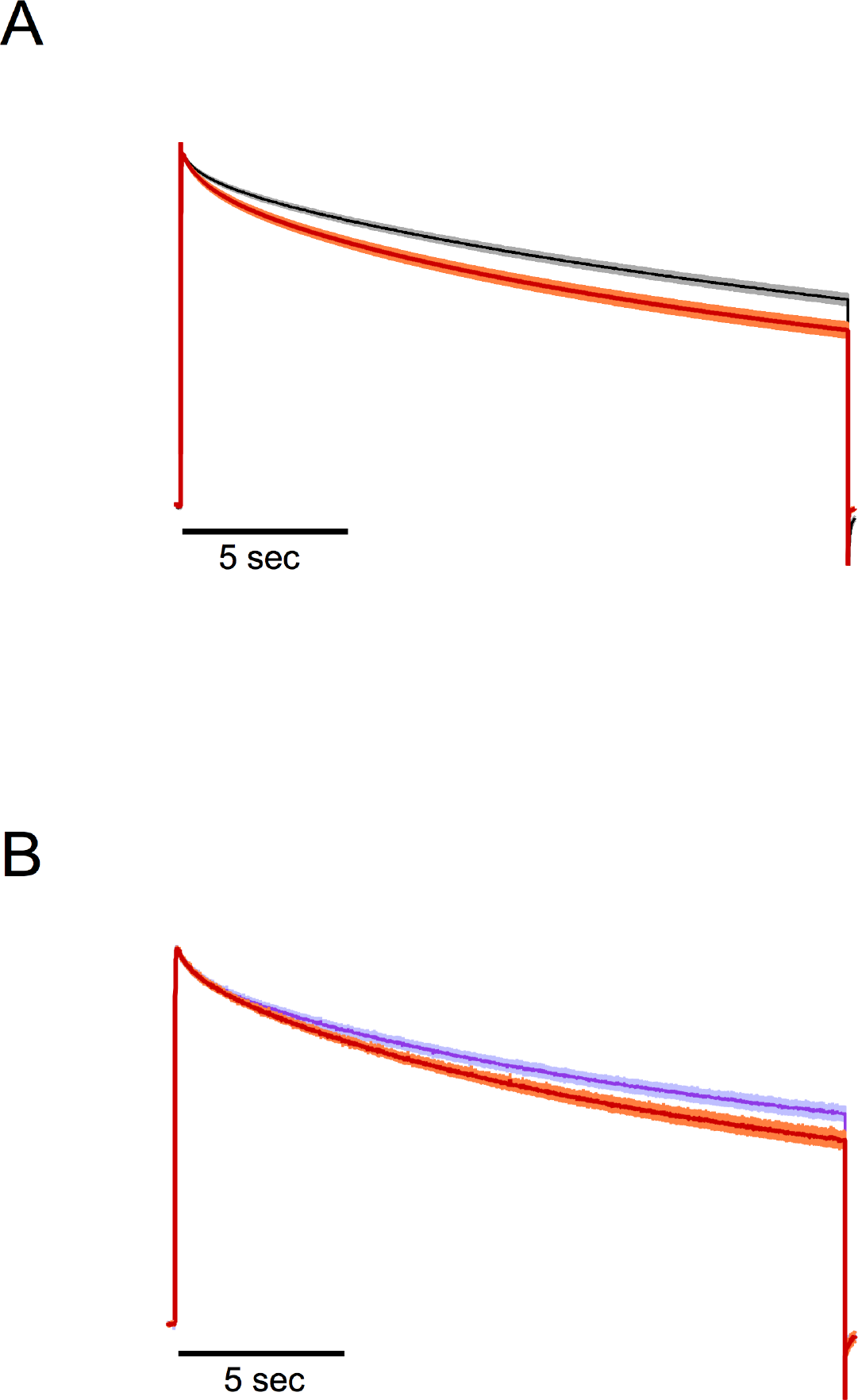
Effect of extracellular potassium and TEA on slow inactivation in WT Kv1.2. (A) Outward currents in response to a 40 mV, 20 s pulse. The red trace is the normalized average current from 7 experiments recorded in 2 mM external K^+^. The black trace is the normalized average current from 9 traces recorded in the presence of 100 mM external K^+^. (B) Effect of external TEA on the kinetics of slow inactivation. The red trace is the normalized average current at 40 mV from 8 traces in the absence of TEA and at 2 mM external potassium. The purple trace is the normalized average current from 9 sweeps in the presence of 20 mM external TEA also with 2 mM external potassium. In A and B, the light colored shade along the current trace is the s.e.m.

In *Shaker* channels, a tryptophan residue at position 434 (W434) has been shown to be very important in modulating C-type inactivation (Perozo et al., 1993; Yang et al., 1997; Pless et al., 2013) and the mutation W434Y has been shown to produce channels with accelerated inactivation (Cordero-Morales et al., 2011). Surprisingly, the equivalent mutant in K_V_1.2, W366Y, produced channels that upon first inspection seem very similar to WT. The channels activate in a voltage-dependent fashion and over a range of voltages almost identical to that of WT (Fig. 3a). Long depolarizing pulses also induce inactivation of the currents, with a double exponential time course that is faster than WTs (Fig 3b). The inactivation process in W366Y is also sensitive to extracellular potassium, but for this mutant it becomes slightly more prominent at [K^+^]_o_ = 100 mM (Fig 3b and c).

**Figure 3.**
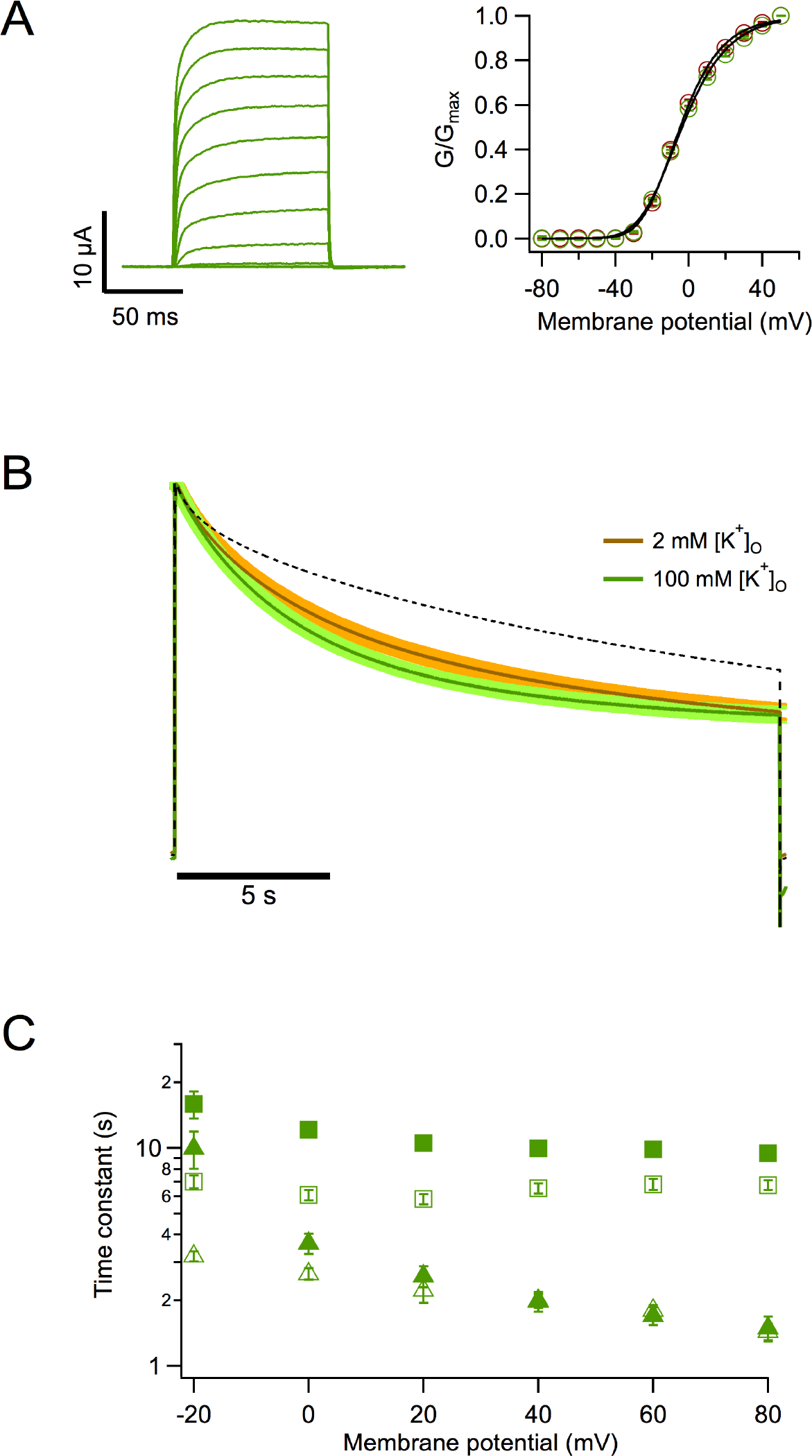
Mutation W366Y accelerates slow inactivation. (A, left) Ionic currents in response to 100 ms depolarizing pulses from −80 mV to 50 mV in 10 mV steps, holding potential was −80 mV. Currents are very similar to WT channels. (Right) Comparison of the voltage dependence of W366Y and WT K_V_1.2. The normalized conductance of both channel types is plotted as a function of voltage. Red circles correspond to WT and green circles to W366Y. The continuous curve corresponds to the fit to Equation 2 with parameters: z_app_ = 1.77 e_o_, K_o_ = 0.135 (WT); z_app_ = 1.59 e_o_, K_o_ = 0.149 (W366Y). (B) Mutant channels also inactivate slowly and more completely than WT channels. High potassium speeds up inactivation. Green trace is the normalized average current at 40 mV from 7 oocytes in the presence of 2 mM external K^+^. Light brown is the normalized current average from 8 oocytes in 100 mN external K^+^. In both cases, the shaded areas are represents ± s.e.m. The black dashed curve is the average WT current time course also at 40 mV. (C) Voltage dependence of the time course of slow inactivation. Filled symbols are the slow and fast time constants in ND96 extracellular medium ([K^+^]_o_ = 2 mM). Empty symbols are the slow and fast time constants in the presence of 100 mM extracellular potassium. Squares are the slow time constant and triangles the fast time constant.

In order to better understand the gating behavior of this mutant, we performed patch-clamp experiments. As seen in TEVC, the ionic currents recorded from inside-out patches of W366Y in response to short depolarizations are similar to WT (Fig. 4a). The slow inactivation elicited by longer pulses (5 s) is also present and occurs along a double exponential time course (Fig. 4b). With the purpose of estimating gating parameters, we performed non-stationary noise analysis experiments. Mean-variance analysis of these currents (Fig. 4c) shows that the single channel current of the mutant is reduced but similar to WT K_V_1.2 (0.53 ± 0.014 pA vs. 0.7 pA at 20 mV, respectively, (Ishida et al., 2015) and that the main effect of the mutation is to significantly reduce the open probability of the channels to near half the value of the WT (Fig. 4d).

**Figure 4.**
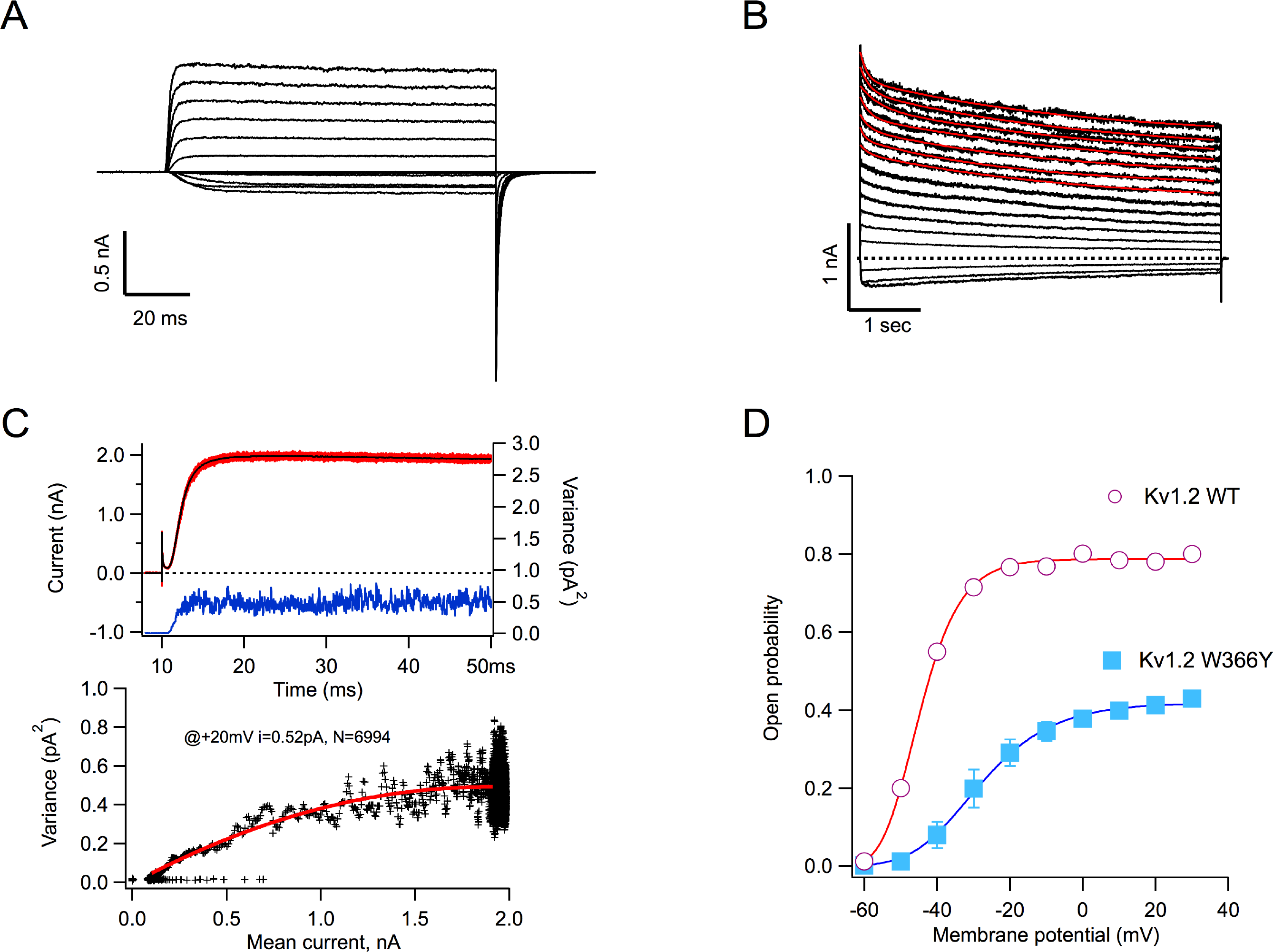
The W366Y mutant induces a rapidly equilibrating state occurring after the open state. (A) Inside-out patch recording of potassium currents through W366Y channels. Currents were elicited by voltage pulses of 100 ms duration, from −60 to 30 mV from a holding potential of −90 mV. (B) Slow inactivation is also present in cell-free recordings and is incomplete at the end of 5 s pulses. The inside-out data show that inactivation proceeds via two exponentials, comparably to the behavior of currents obtained in TEVC (red curves are the fits to a double-exponential function). (C) Non-stationary noise analysis of currents at 20 mV. In the upper panel one hundred current sweeps are shown in red with the mean current plotted in black. The blue trace is the point-by-point variance calculated from the 100 traces. The bottom panel shows the mean-variance relationship (black crosses) with the fit to Equation 1 shown in red. The parameters of the fit are: I = 0.52 pA, N = 6994. (D) The voltage dependence of activation of WT (red circles) and W366Y (blue squares) channels. The amplitude of the tail current at the end of 100 ms pulses is plotted as a function of the voltage of the pulse and is normalized to the maximum open probability obtained from noise measurements as in (C). Data are mean ± s.e.m. WT, n = 10; W366Y, n = 7.

Another mutation in *Shaker* that is known to modulate the time course of C-type inactivation is the W434F substitution, which accelerates inactivation to such extent that channels with extremely reduced open probability and transient openings are produced. As a consequence, when enough channel expression is achieved, only charge movement in the form of gating currents is observed, as if channels were permanently C-type inactivated (Perozo et al., 1993; Yang et al., 1997).

Surprisingly, mutation W366F, which is equivalent to *Shaker* W434F, only produces rapidly inactivating channels (Fig. 5a). This result had been previously reported (Cordero-Morales et al., 2011). Side by side comparison of normalized currents of the two mutants at position 366 and WT is shown in Figure 5b. It is readily seen that W366F channels inactivate within milliseconds, with a time course that can be fit with a single exponential. This time-constant is 2.5 orders of magnitude faster than the fastest component of inactivation in WT or W366Y (Fig. 5c). This fast inactivation was also observed in inside-out patch recordings (Fig. 5d). These inside-out recordings were performed in the presence of 60 mM [K^+^]_o_. Under these conditions, the conductance activates at more negative voltages, as compared with the conductance in the presence of 2 mM [K^+^]_o_, also in inside-out recordings (Fig. 5e).

**Figure 5.**
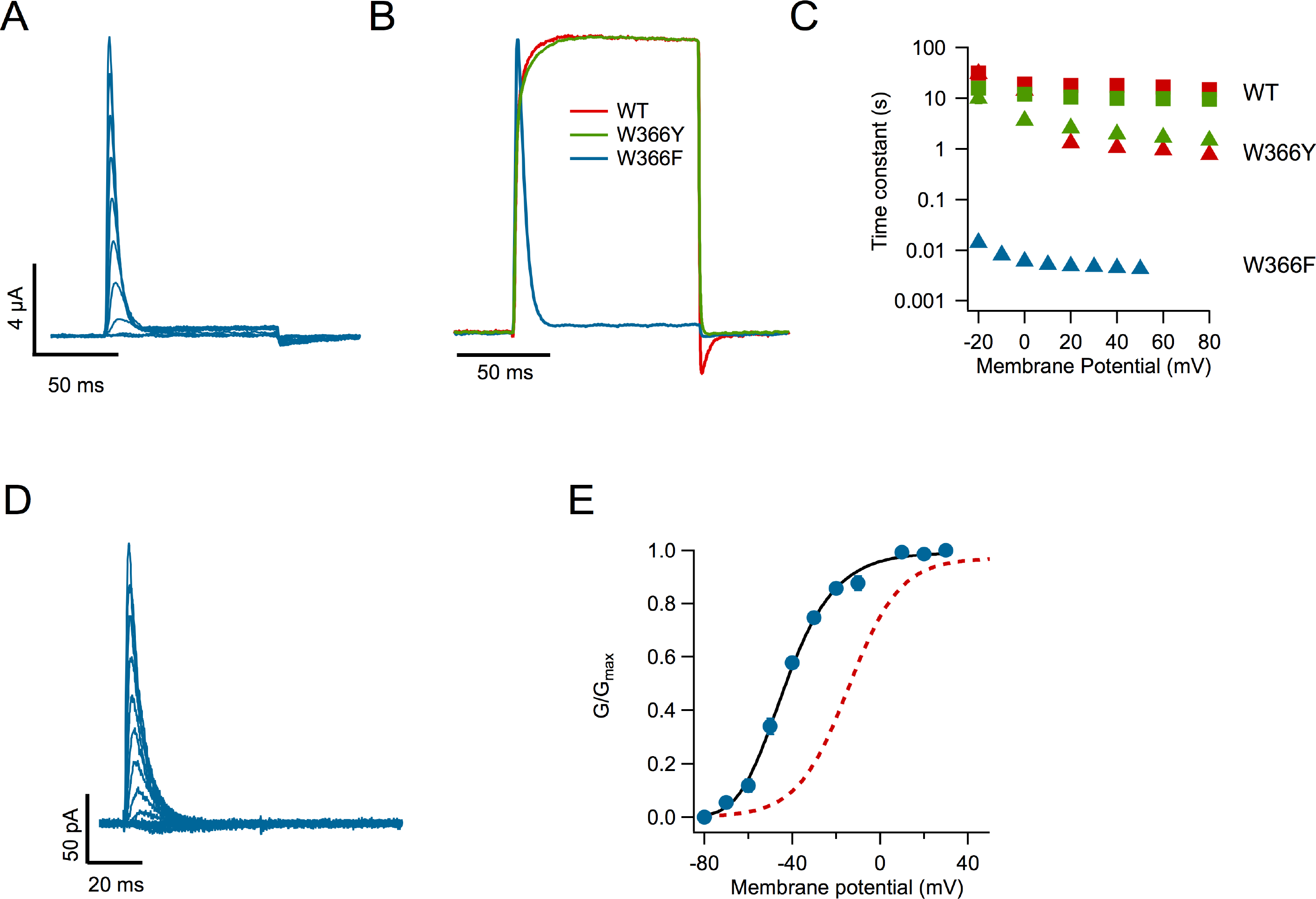
The W366F mutation produces channels with fast inactivation. (A) Family of currents showing fast inactivation even with short (100 ms) pulses. Depolarizing pulses stepped from −80 mV to 50 mV in 10 mV increments, holding potential was −80 mV (B) Comparison of the time course of currents activated by a 100 ms pulse to 50 mV for WT, W366F and W366Y channels. Currents were normalized to their maximum value. (C) Inactivation time constants for the two mutants and WT channels. The fast and slow time constants for WT and W366Y were obtained from currents elicited by 20 s pulses as in figures 1d and 3c. The inactivation time constant for W366F was obtained from a fit to a single exponential function of the decay of currents elicited by 100 ms. (D) Fast-inactivating currents obtained from an inside-out patch expressing W366F channels. (E) The voltage dependence of these currents is shifted to negative voltages when 60 mM extracellular potassium is used. The continuous black curve is the fit to equation 2 with parameters z_app_ = 1.75 e_o_, K_o_ = 0.0091 (n = 6). For comparison, the voltage dependence of currents in 2 mM external potassium is shown (red dotted curve).

An important question to ask is if the inactivated state that seems to be stabilized in K_V_1.2-W366F is the same slow-inactivated state seen in WT and W366Y. As with these channels, application of 100 mM extracellular K^+^ to K_V_1.2-W366F results in a significant slowing of the inactivation rate (Fig. 6a and b), as reported for C-type inactivation in other potassium channels. The extent of this slowing is larger than in WT, reaching ~3.4-fold vs. 1.3 at positive voltages.

**Figure 6.**
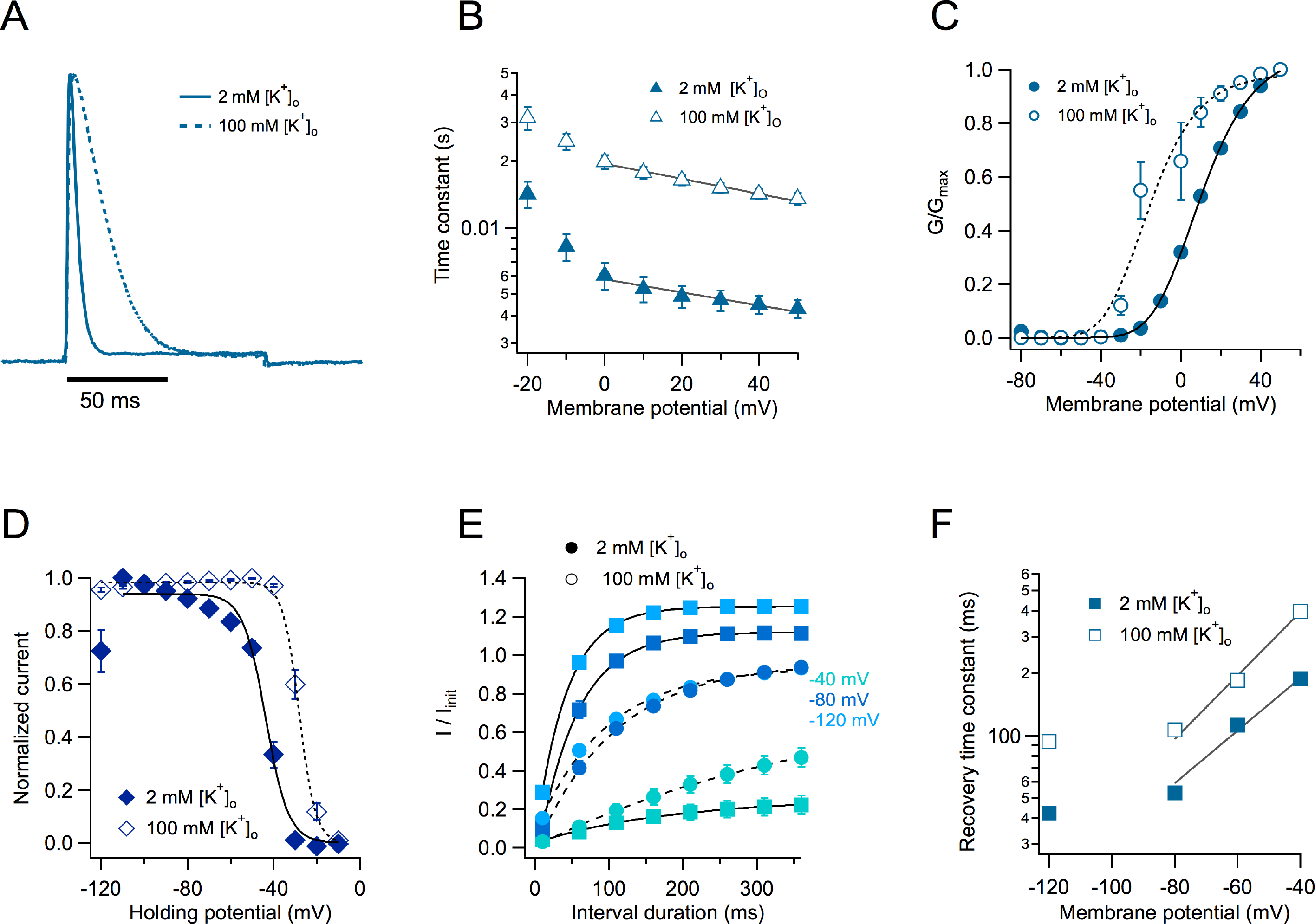
Inactivation characteristics of the W366F mutant. (A) Current elicited by a 50 mV pulse of 100 ms duration obtained in the presence of 2 mM extracellular K^+^ (continuous line). The dotted line is a representative current at the same voltage in the presence of 100 mM extracellular K^+^. (B) The inactivation time constant obtained from a single exponential fit to currents in the presence of 2 mM [K^+^]_o_ (filled symbols) or 100 mM [K^+^]_o_ (hollow symbols). The black lines show an exponential fit to equation 6. The apparent charges in high and low external potassium are similar (−0.17 and −0.19 e_o_, respectively) and the value of τ_o_ is increased from 0.005 s to 0.019 s by the high potassium. (C) Effect of external potassium on the voltage dependence of the peak current. Filled symbols are the normalized peak conductance in 2 mM [K^+^]_o_ from n = 9 experiments and the continuous line is the fit to equation 2 with parameters, z_app_ = 1.52 e_o_, K_o_ = 0.35. Hollow symbols are the normalized peak conductance in 100 mM [K^+^]_o_. The parameters of the fit (dashed curve) are: z_app_ = 1.64 e_o_, K_o_ = 0.068 (n = 9). (D) Steady state inactivation. A prepulse of varying voltage from −120 mV to −10 mV of 300 ms duration was applied before a fixed 100 ms pulse to 50 mV. The current elicited by the 50 mV pulse was normalized and plotted as a function of prepulse voltage. The data is fitted by equation 4. Filled symbols are in 2 mM [K^+^]_o_ and hollow symbols in 100 mM [K^+^]_o_. The parameters of the fit are: 2 mM [K^+^]_o_ and 100 mM [K^+^]_o_. (E) Recovery from inactivation was measured with a two pulse protocol with pulses of 100 ms duration at 50 mV. The pulses were separated by increasing time intervals from 10 to 360 ms in 50 ms increments. The voltage of the interval between two pulses was held at −120, −80 and −40 mV. The recovery time course was fitted to a single exponential function and the voltage dependence of the recovery was plotted in F. The experiment was done at 2 mM [K^+^]_o_ (filled symbols, n = 9) and 100 mM [K^+^]_o_ (hollow symbols, n = 9). Mean and s.e.m. are plotted. (F) Recovery rates are shown as a function of prepulse voltage and their voltage dependence was assessed by a fit to equation 6, indicating that the apparent charge of recovery is increased by 100 mM external K^+^ from 0.73 e_o_ to 0.88 e_o_.

As shown in inside-out patch clamp recordings (Fig. 5e), in TEVC increased [K^+^]_o_ also modulates the range of activation by voltage. 100 mM [K^+^]_o_ shifts the mid point of activation by −30 mV (Fig. 6c). The antagonistic action of elevated [K^+^]_o_ on inactivation is also evidenced by the effects on the voltage dependence of steady-state inactivation. High [K^+^]_o_ shifts the voltage dependence of inactivation by +20 mV. Importantly, a significant proportion of channels (26 %) is inactivated at −120 mV in 2 mM [K^+^]_o_. This inactivated fraction of channels at negative voltages disappears when [K^+^]_o_ is brought to 100 mM (Fig. 6d).

Recovery from inactivation is highly voltage–dependent, becoming faster as the holding potential is made more negative (Fig. 6e) and the rate of recovery is also modulated by [K^+^]_o_. However, the modulation of inactivation by external potassium is counterintuitive, since 100 mM [K^+^]_o_ makes recovery slower and not faster, as would be expected from potassium’s slowing of the rate of entering the inactivated state (Fig. 6f).

Extracellular TEA has also been used as a probe of C-type inactivation (Kurata et al., 2005; Carrillo et al., 2013). As with WT K_V_1.2 (Fig. 2), we applied 20 mM TEA to the bath and observed a modest but robust slowing of the inactivation rate of K_V_1.2-W366F (Fig. 7a and b), a result consistent with similar experiments in *Shaker* channels. Remarkably, application of intracellular TEA (20 mM) to inside-out patches, also slowed down inactivation, although the effect was visible at more negative voltages that with extracellular TEA (Fig 7c).

**Figure 7.**
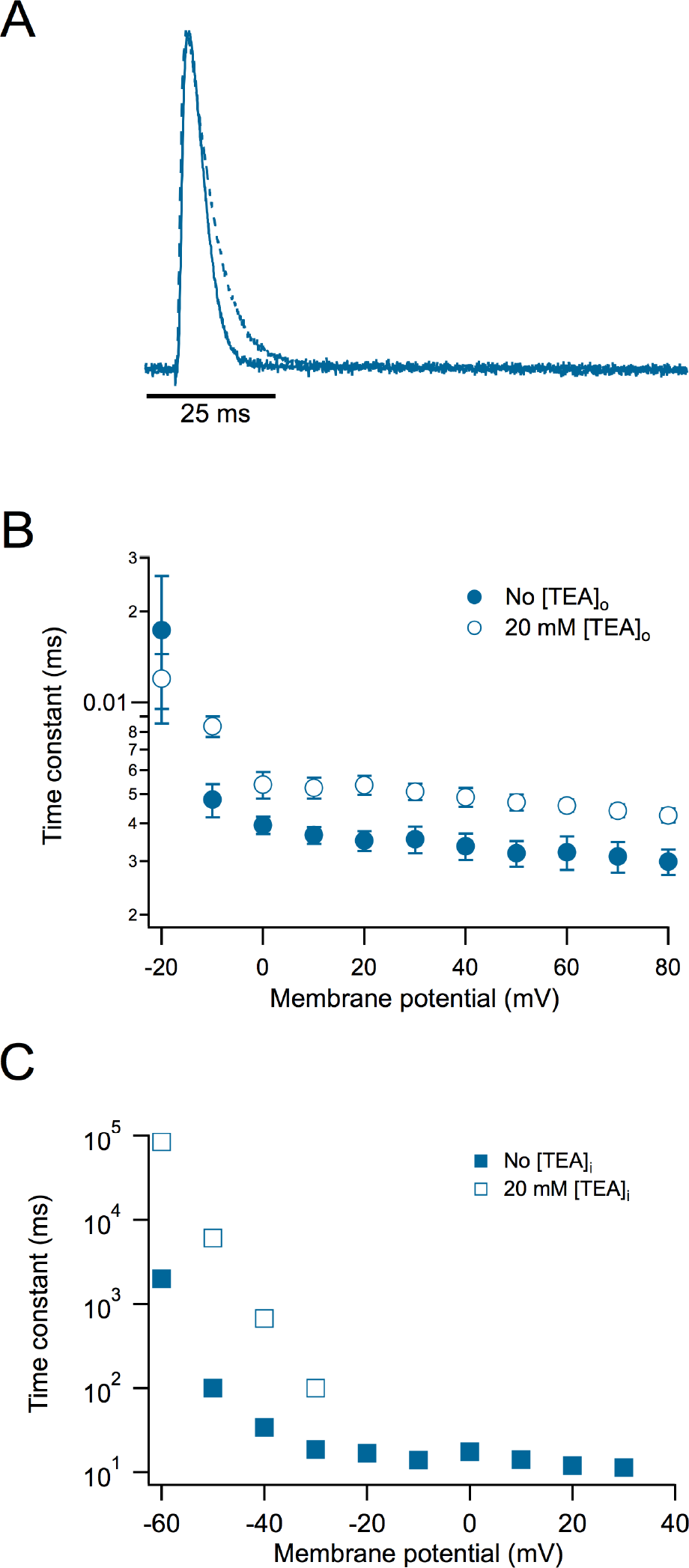
Extracellular and intracellular TEA slows down the inactivation kinetics of the W366F mutant. (A) Representative current traces at 60 mV in the absence (continuous trace) and presence of 20 mM extracellular TEA (dashed trace). (B) Voltage dependence of the time constant of inactivation. External TEA data is show by hollow symbols, while control data is shown by the filled symbols. (C) Effect of intracellular TEA. 20 mM intracellular TEA was applied to inside-out patches. In the presence of TEA, current inactivation is slowed down appreciably at negative voltages. Data shown are mean and s.e.m.

Taken together, these set of experiments with [K^+^]_o_ and [TEA]_o_, suggest that the inactivation induced by the K_V_1.2-W366F mutant has characteristics compatible with it being an accelerated form of the slow inactivation present in WT channels.

Why is it that the W366F mutation in K_V_1.2 does not produce permanently inactivated channels as in *Shaker*? Several amino acid residues have been shown to be involved in modulating the rate of slow inactivation in *Shaker* channels. In particular, when a threonine is located at position 449 (*Shaker* WT) the channels inactivate in the course of seconds. Inactivation is greatly enhanced by a charged (positive or negative) amino acid or an alanine, while valine almost completely abolishes inactivation (López-Barneo et al., 1993). In K_V_1.2 channels the equivalent position is V381, and when mutated to threonine, combined into the double mutant W366F/V381T (Goodchild et al., 2012) channels that only display gating currents are produced.

We also made the W366F/V381T double mutant, along with W366F/V381A, W366F/V381I, W366F/V381W, W366F/V381L. Of these double mutants, only W366F/V381A, W366F/V381I and W366F/V381T expressed functionally. These three double mutants gave rise to channels that only displayed charge movement. Gating currents were readily recorded in whole oocytes (Fig. 8a) and showed characteristics that are similar to WT gating currents recorded in oocytes via patch-clamp (Ishida et al., 2015). The Q-V relationship shows that charge moves in two main components (Fig. 8b) that occur at voltages more negative than the peak conductance of the background mutant, W366F. The kinetics of the on-gating currents is similar between mutants, especially at positive potentials. At negative voltages, the mutant W366/V381T shows slightly slower kinetics. The off-gating currents are variously contaminated by the tail currents of endogenous oocyte channels, so a comparison of on-charge vs. off-charge was not carried out.

**Figure 8.**
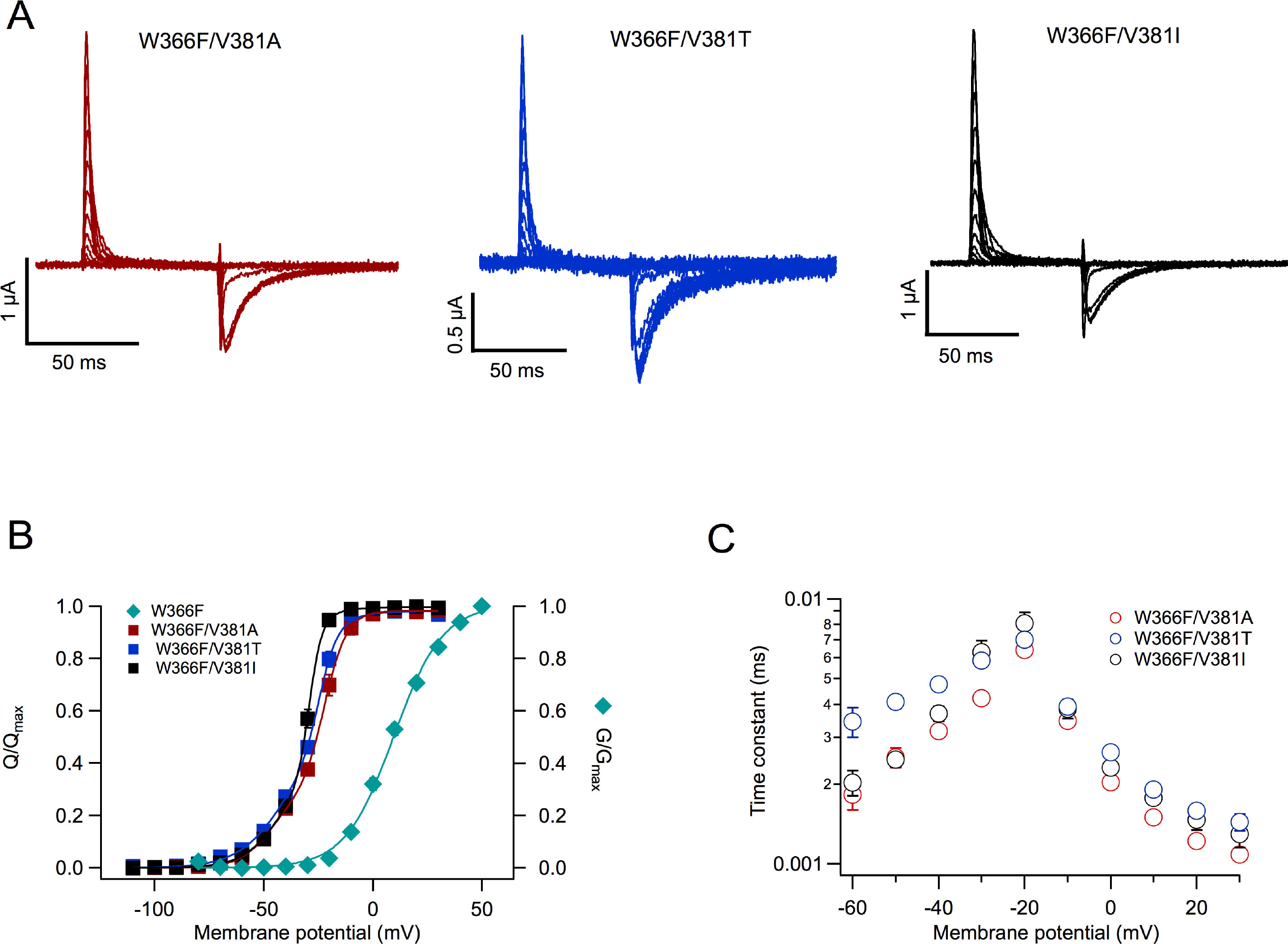
Mutations at V381 in the W366F background produce permanently inactivated channels. (A) Representative traces of gating currents recorded from the expressed channels indicated in each panel. Currents were elicited by voltage-clamp pulses of 60 ms duration ranging from −110 to 30 mV in 10 mV steps from a holding potential of −100 mV. (B) Normalized voltage dependence of charge movement (Q-V) of the three mutants plotted (following its color code from A) and compared to the voltage dependence of the peak-normalized conductance of the background mutation W366F (cyan rhombuses). Q-V data was fitted to equation 5. The number of experiments are: W366F/V381A, n = 13; W366F/V381T, n = 5; W366F/V381I, n = 9. (C) The time course of current decay of the three double mutants is plotted as a function of pulse voltage. On-gating current decay was fitted to a single exponential to determine the time constant. Number of experiments is as in B. Data are mean and s.e.m.

## Discussion

Slow inactivation in potassium channels is a gating transition of functional importance. In spite of this, its fundamental mechanism remains not well understood. This is in part due to the fact that more than one form of inactivation coexists in several channels. Even the better understood *Shaker* channel shows evidence of more than one way to enter an inactivated state (Ayer Jr and Sigworth, 1997; Klemic et al., 2001). C-type inactivation (sometimes also called C/P-type (Kurata and Fedida, 2006); (Bähring et al., 2012a) seems to be associated with conformational changes in the selectivity filter that occur after the channel has reached the open state (Scheme I). U-type inactivation seems to be related to conformational changes in the pore that can occur while the channel is in its closed conformation (Bähring and Covarrubias, 2011). These two forms of inactivation coexist in *Shaker* channels and constitute an often overlooked confounding factor in inactivation studies.

Here we have shown that WT K_V_1.2 channels slow-inactivate and this inactivation process also has mixed characteristics of C/P-type and U-type inactivation. Compared to slow inactivation in *Shaker* (Hoshi et al., 1991), K_V_1.2’s is less pronounced. This is perhaps due to the presence of a valine at position 382, which in *Shaker* is a threonine. Valine at the equivalent position in *Shaker* also produces channels with less complete and slower inactivation (López-Barneo et al., 1993).

A simplified kinetic model can explain the behavior of WT and the mutations studied in this work (Scheme I).

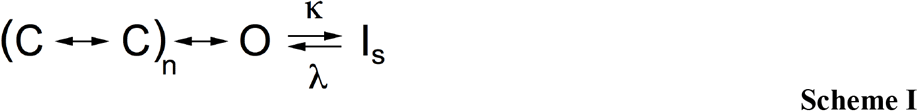

A model like Scheme I assumes sequential charge movement and a single open state. Slow inactivation (I_s_) can only occur from the open state. For the W366Y channels, the rate constant κ is increased with respect to WT, leading to faster and more complete inactivation at the end of a long activating pulse. Additionally, the rate constants previous to the open state are also reduced by the mutation W366Y, since the peak open probability is reduced in this mutant.

The behavior of the W366F mutant can be explained if κ is much greater than λ, leading to a much faster and complete inactivation. This model is not unique and its likely only part of a full explanation of the mutants’ effects. Since WT K_V_1.2 shows evidence of U-type inactivation, this would require the existence of inactivated states accessed from closed states (Cheng et al., 2011; Bähring et al., 2012b; Jamieson and Jones, 2013).

Evidence that slow inactivation in WT and the mutants is similar to C-type inactivation in *Shaker*, is provided by the effects of external K^+^ and TEA (Levy and Deutsch, 1996). Both compounds slow down the time course of inactivation in WT and the W366Y and W366F mutants, which is indicative of a common mechanism.

The behavior of W366F is very interesting. These channels have a fast and complete inactivation that is not very voltage-dependent (0.17 e_o_) and becomes slower with high extracellular potassium, but not significantly different in voltage dependence (0.19 e_o_). However, recovery from inactivation is more voltage-dependent (0.73 e_o_) and becomes even more so in high external potassium (0.88 e_o_). External potassium has a paradoxical effect on inactivation. While it slows down entry into the inactivated state and removes the steady-state inactivation present at negative voltages, it also slows down recovery from inactivation.

The effects of extracellular TEA have been used as a probe of the conformational changes occurring in the external region of the pore during slow inactivation (Andalib et al., 2004). We find that as in *Shaker*, application of external TEA slows down the transition to the inactivated state. This is indicative of similar steric and electrostatic interactions in both channels during inactivation.

It has been show that in *Shaker*, intracellular TEA interferes with C-type inactivation more prominently than external TEA (González-Pérez et al., 2008), suggesting that internal TEA might hinder the U-type inactivation component while external TEA slows down C-type inactivation. Surprisingly, we find a similar behavior in W366F, where application of intracellular TEA to this mutant also slowed down inactivation rate. Our experimental evidence suggests that the conformational changes leading to a slow-inactivated state in K_V_1.2 and *Shaker* are similar. However the detailed effects of the mutants are not the same in these two channels.

The phenotype of W366F is remarkable in that it does not produce permanently C-type inactivated channels. It was previously demonstrated that this mutation combined with V381T, which mimics the background of the mutant W434F in *Shaker*, abolishes ionic current and produces channels with only gating currents (Goodchild et al., 2012).

When other amino acid residues are substituted at position V381 in the W366F background, channels with only gating currents are also produced. Of all the mutants we tried, only W366F/V381A, W366F/V381I and W366F/V381T produced functional channels. A bulkier amino acid such as tryptophane was not tolerated, as was leucine instead of isoleucine. We believe that the effects of the mutants can be interpreted in terms of a mechanism that might be responsible for the initiation and establishment of the slow-inactivated state.

### A sequence of events leading to slow inactivation

The effects of the mutants presented in this work lead us to propose a sequence of events that in conjunction constitute slow inactivation in K_V_1.2 channels (Fig. 9). Prolonged flux of potassium through the selectivity filter leads to a conformational change (perhaps initiated by strong electrostatic interactions) that starts at tyrosine Y377. Rotation of this tyrosine into W366, which is initially between 3 and 4 Å apart, will destabilize the interaction of a K^+^ ion with the S2 binding site in the selectivity filter. In turn, the hydrogen bond between W366 and D379 (Pless et al., 2013) will be destabilized, leading to further conformational changes in the external part of the channel pore. In the presence of a threonine at position 381, as in *Shaker* channels, this destabilization will be enhanced due to a repulsive dipole-charge interaction.

**Figure 9.**
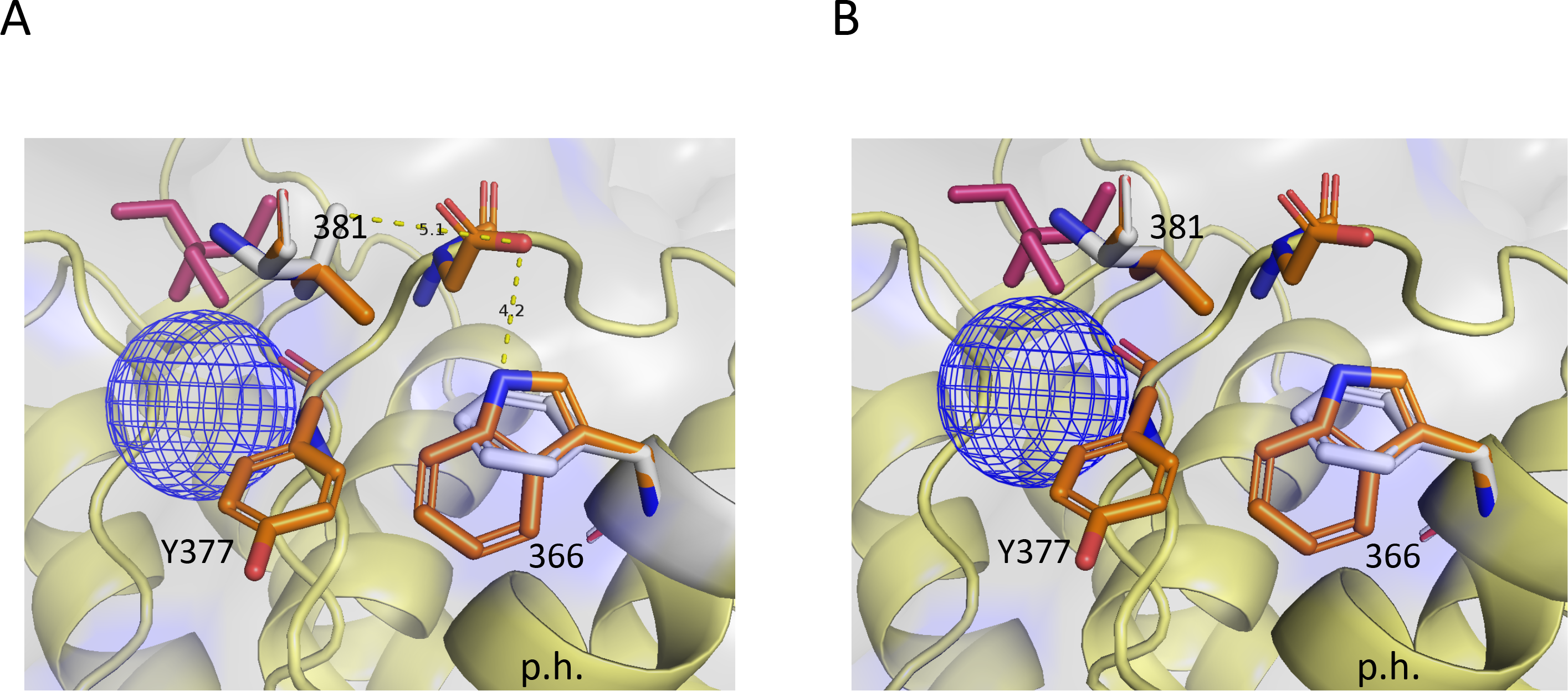
Structural basis of modulation of inactivation by mutants at positions W366 and V381. (A) Residues involved in slow inactivation near the extracellular region of the selectivity filter of the Kv1.2-Kv2.1. The stick representation in colors corresponds to residues W366, V381, Y377 and D379. The white, superimposed residues are the changes introduced by the double mutations W366F/V381T. The main changes in this mutant are: an increased space and reduced interaction between W366F and Y377 and elimination of the hydrogen bond between W366 and D379 (dotted yellow line). V381T might enhance the effects of W366F because it should repel D379. (B) The double mutant W366F/V381A. Residue changes are again represented in white. In this mutant there is no interaction between positions 381 and D379. The blue mesh sphere is the density of a potassium ion occupying the S1 site in the selectivity filter. The purple stick model is a TEA molecule occupying the S0 site. High occupancy of these two sites might antagonize slow inactivation through steric hindrance. The legend p.h. indicates the pore helix.

When a tyrosine or a phenylalanine is present at 366, there is more space for Y377 and no hydrogen bond to be formed with D379, so the inactivated state is formed with faster kinetics. In the double mutant W366F/V381T this inactivation is enhanced due to repulsive interactions between T381 and D379. The mutants W366F/V381A, W366F/V381I must produce similar end effect but through different mechanism; the presence of A381 leaves so much empty space and no interactions with D379 that the entire conformation of the pore is altered. Similarly, I381 possibly leaves little space and pushes D379 away.

Overall, there are multiple inter and intra-molecular interactions leading to the slow inactivated state. The mutants that we have analyzed in K_V_1.2 might be involved only in the conformational changes in the external parts of the pore. If U-type inactivation is part of the slow-inactivated state, these separate conformational changes might occur in the inner pore vestibule, as has been previously suggested (González-Pérez et al., 2008).

## Acknowledgments

This work was financed by grants from CONACyT (252644) and UNAM-DGAPA-PAPIIT (IN209515) to LDI. We thank Itzel Llorente from Instituto de Fisiología Celular, UNAM for technical support. The authors declare no competing financial interests.

## Author contributions

Esteban Suarez-Delgado carried out electrophysiology experiments, data analysis and read the manuscript. Teriws G. Rangel-Sandín, carried out electrophysiology experiments and read the manuscript. Itzel G. Ishida carried out patch-clamp experiments, data analysis and contributed to making mutants. Gisela E. Rangel-Yescas contributed reagens, carried out molecular biology experiments and read the paper. Tamara Rosenbaum, conceived research, wrote and read the paper. León D. Islas conceived research, carried out data analysis, contributed analysis programs and wrote the paper.

